# Outer membrane permeabilization by the membrane attack complex sensitizes Gram-negative bacteria to antimicrobial proteins in serum and phagocytes

**DOI:** 10.1101/2020.01.28.923243

**Authors:** Dani A. C. Heesterbeek, Remy M. Muts, Vincent P. van Hensbergen, Pieter de Saint Aulaire, Tom Wennekes, Bart W. Bardoel, Nina M. van Sorge, Suzan H.M. Rooijakkers

## Abstract

Infections with Gram-negative bacteria form an increasing risk for human health, which is mainly due to the increase in antibiotic resistance. The cell envelope of Gram-negative bacteria consists of an inner membrane, a peptidoglycan layer and an outer membrane, which forms an impermeable barrier to many antibiotics and antimicrobial proteins. The complement system is an important factor of the human immune system that can efficiently kill Gram-negative bacteria via the formation of large pores in the bacterial outer membrane, called Membrane Attack Complexes (MACs). To better understand how these MAC pores damage the complex cell envelope of Gram-negative bacteria, we recently developed a fluorescent reporter system to study membrane damage in *E. coli*. Here, we used a similar experimental setup in combination with flow cytometry, confocal microscopy and conventional plating assays to elucidate how different components of the immune system act synergistically to effectively clear invading bacteria. We demonstrate how MAC-dependent outer membrane damage enhances the susceptibility of *E. coli* to further degradation of the cell envelope by lysozyme, leading to drastic changes in the morphology of these bacteria. Furthermore, we elucidate that the MAC enhances the susceptibility of *E. coli* to phospholipases, and to degradation and killing inside human neutrophils. Altogether, this study provides a detailed overview on how different players of the human immune system work closely together to degrade the complex cell envelope of Gram-negative bacteria. This knowledge may facilitate the development of new antimicrobials that could stimulate, or work synergistically with the immune system.

## Introduction

Infections with Gram-negative bacteria form a major problem for human health, which is mainly due to the increase in antibiotic resistance. According to the World Health Organization there is an urgent need for alternative strategies to treat infections with Gram-negative bacteria such as *Acinetobacter baumanni, Pseudomonas aeruginosa* and *Escherichia coli (E.coli)*, which are at the top of the priority list of antibiotic resistant bacterial species^1^. The cell envelope of Gram-negative bacteria consists of an inner membrane, a peptidoglycan layer and an additional outer membrane^2^. The peptidoglycan layer of bacteria is important for the maintenance of osmotic balance and to maintain the bacterial cell shape, for example rod-shaped for *E. coli*^3^. The outer membrane forms a physical barrier to a large number of antimicrobial compounds^4^, which makes it challenging to develop new antibiotics against these bacteria. Combination therapy of antibiotics and outer membrane permeabilizing agents have become more attractive over the last decades^5–9^. Furthermore, there is increased awareness that antibiotics may be more effective in the presence of the human immune system^10^, which has developed strategies to disrupt the complex cell envelope of Gram-negative bacteria. Increased understanding of these mechanisms may facilitate the development of new antimicrobials, that could stimulate, or work synergistically with the immune system.

The human body fights invading bacteria via cellular and humoral immune components. Cellular protection is provided by immune cells, such as neutrophils, that engulf bacteria and expose them to a large number of antimicrobial compounds^11,12^. One of these intracellular proteins is lysozyme, a 14.7 kDa protein that degrades bacterial peptidoglycan^12,13^. Besides being present in immune cells, lysozyme is also present in body fluids such as the blood, saliva and tears^13^. Although lysozyme and other (intracellular) antimicrobials can efficiently act on Gram-positive bacteria^14^, many of these are inactive against Gram-negative bacteria^15,16^, partly because they cannot cross the bacterial outer membrane. Humoral immunity against bacteria is mainly dependent on the complement system, which consists of a protein network circulating in the blood. Complement activation triggers the deposition of C5 convertases on the bacterial surface that cleave C5 into C5a and C5b. C5b associates with C6, C7, C8 and multiple copies of C9 to form large ring-structured pores (C5b-9)^17,18^, called Membrane Attack Complexes (MACs). It has long remained unclear how MAC pores damage the complex cell envelope of Gram-negative bacteria in such a way that this leads to bacterial cell death. Furthermore, although the bactericidal activity of the MAC has been analyzed in combination with other immune factors^19,20^, tools to study these processes at a molecular level were limiting.

We recently developed a fluorescent reporter system to distinguish between outer and inner membrane perforation in Gram-negative bacteria by flow cytometry. Specifically, *E. coli* was genetically engineered to express mCherry in the periplasm and GFP in the cytoplasmic space. Leakage of these proteins and influx of impermeable DNA dyes functioned as a detailed readout for membrane damage. Using these tools, we demonstrated that the MAC efficiently permeabilizes the bacterial outer membrane which can, after a delay, also trigger destabilization of the bacterial inner membrane^10,18^. Whereas outer membrane damage by itself is not sufficient to kill Gram-negative bacteria, inner membrane damage is crucial to prevent colony formation. By studying how MAC pores affect membrane integrity using purified complement proteins, we also noticed that the MAC can efficiently kill Gram-negative bacteria, but does not trigger leakage of cytoplasmic proteins (GFP). Furthermore, MAC pores do not alter the cell shape of bacteria^10,18^, suggesting that the peptidoglycan layer remains intact. This suggests that although the MAC is able to directly kill Gram-negative bacteria, other factors are required to further degrade the bacterial cell envelope.

Here we demonstrate, at a molecular level, how MAC-dependent outer membrane damage enhances the susceptibility of *E. coli* to antimicrobial proteins with different effector functions, such as human lysozyme and Group IIA secreted phospholipase A2 (hGIIA)^21^. Lysozyme turned out to be the crucial factor for the disintegration of the cell wall of *E. coli* in serum. In addition, we show that MAC-dependent outer membrane damage enhances killing and degradation of bacteria inside human neutrophils, suggesting that it sensitizes bacteria to antimicrobial proteins in the phagolysosome. Altogether, this study describes how different components of the immune system act synergistically to effectively clear invading bacteria.

## Results

### The MAC and lysozyme trigger E. coli cell wall degradation in human serum

In previous studies, we investigated how the MAC affects the cell wall integrity and viability of *E. coli* using purified complement components. We observed that the MAC efficiently damages the outer and inner membrane of bacteria, which triggers bacterial cell death. Nevertheless, confocal microscopy images revealed that MAC-treated bacteria remain rod-shaped, which is in line with the fact that these bacteria retained their forward and side scatter when measured by flow cytometry^10,18^. In contrast to these purified conditions, we noticed that *E. coli* bacteria start to disappear from the flow cytometry gate after approximately 40-60 minutes when these are incubated with full serum (data not shown). To investigate this further, we adjusted the acquisition settings of our flow cytometry experiment to quantify the number of particles in a sample, by measuring a fixed volume. Particles were qualified as rod-shaped bacteria when their forward scatter (FSC) and side scatter (SSC) was similar to that of untreated bacteria (**Fig. 1a**). A threshold was set on the SSC, to prevent the detection of background events caused by serum or by bacteria that have a different morphology. Exposing bacteria to serum drastically decreased the number of detected particles, suggesting that these bacteria lost their natural rod-shape (**Fig. 1a, b**). Since the bacterial peptidoglycan layer is important in maintaining the bacterial cell shape and to prevent bacterial lysis from turgor pressure^22^, we questioned whether lysozyme in serum could be responsible for the observed loss of particles. Although lysozyme is normally not active against *E. coli*, we hypothesized that the combined action of complement and lysozyme in serum might promote the degradation of the bacterial cell wall. To test this, we exposed bacteria to serum from which >99% of the endogenous lysozyme was depleted (Δlysozyme serum)^18^. In the absence of lysozyme, bacteria retained the same FSC/SSC as untreated bacteria, suggesting that these bacteria remained rod-shaped. The effect of non-depleted serum was restored when lysozyme was added back to the Δlysozyme serum (**Fig. 1a, b**). The repleted serum was even more efficient than the non-depleted serum, likely because the added lysozyme concentration exceeds that of normal serum with ∼100x. The efficacy of lysozyme in serum was depended on the presence of MAC pores, as no particles disappeared in the presence of the C5 inhibitor OmCI, which prevents MAC formation^23^ (**SFig. 1a**).

**Figure 1:**
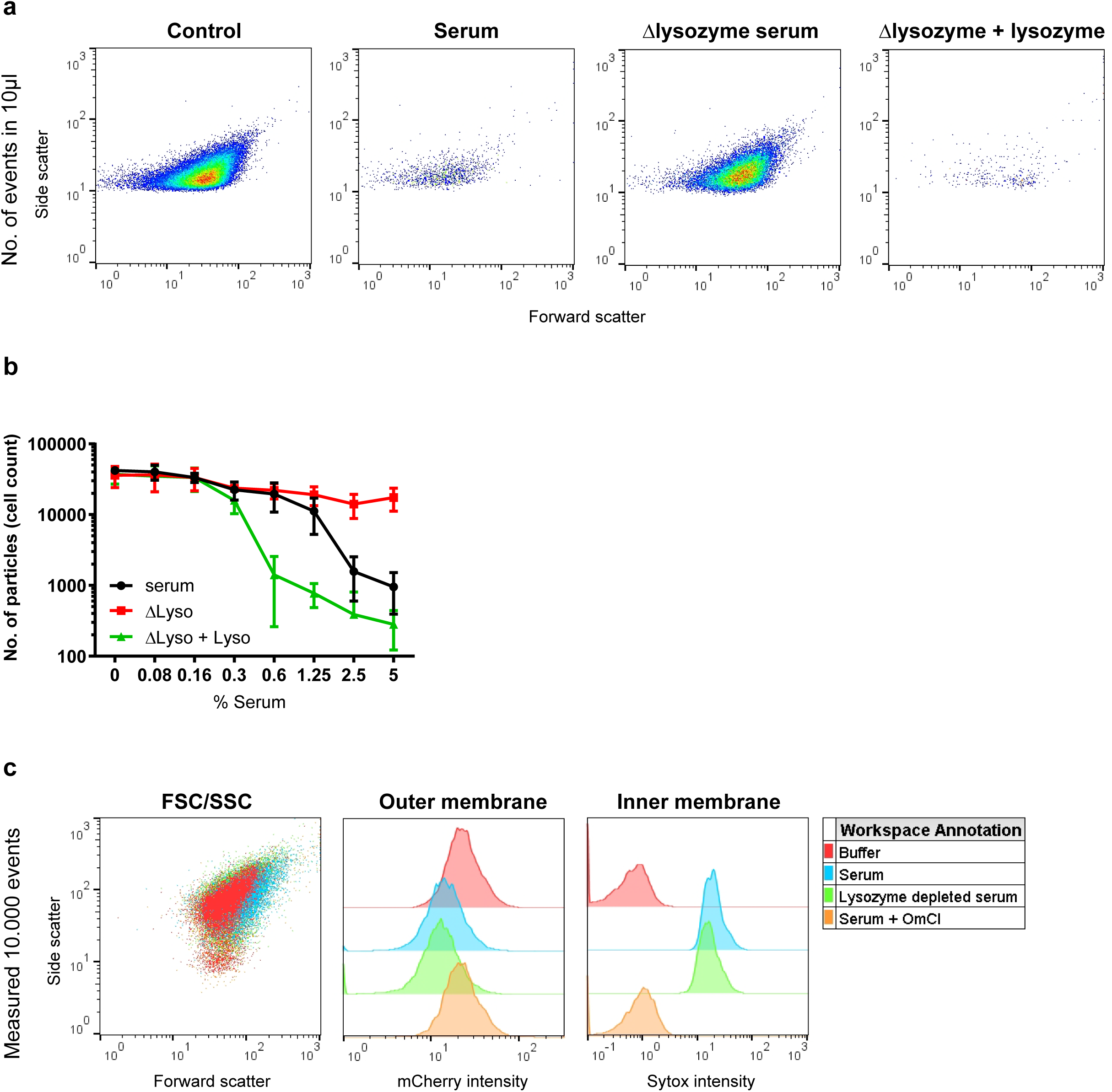
The MAC and lysozyme trigger *E. coli* cell wall degradation in human serum. **A**) Flow cytometry plots (FSC/SSC) of the number of *E. coli* particles in 10 µl after exposure to buffer, 5% serum or Δlysozyme serum with or without 5 µg/ml lysozyme for 60 minutes at 37°C. A SSC threshold was set based on untreated bacteria to filter out background noise and to determine changes in FSC/SSC upon treatment with different serum conditions. **B**) Cell count in 10 µl incubation volume of *E. coli* treated with a concentration range of nhs or Δlysozyme serum with or without 5 µg/ml lysozyme for 60 min 37°C. The number of cells represent the events in 10 µl sample that were measured within the conditions depicted in **A. C**) Flow cytometry plot (left, FSC/SSC) and histograms of outer membrane damage (middle: mCherry) and inner membrane damage (right: Sytox blue) of bacteria treated with buffer, 1% serum with or without 20 µg/ml OmCI or 1% Δlysozyme serum. **A, C**) Flow cytometry plots and histograms represent data of three independent experiments. **B**) Data represent mean ±SD of 3 independent experiments.

Since we observed that lysozyme plays a critical role in serum-mediated cell wall degradation, we questioned whether lysozyme contributes to outer or inner membrane damage. Although we previously observed that lysozyme is not essential for inner membrane damage or killing of *E. coli* in 10% serum^18^, it may play a more crucial role in lower serum concentrations. To study potential damage to each of the two bacterial membranes separately, we used the genetically engineered *E. coli* strain that expresses mCherry in the periplasm and GFP in the cytoplasm (_peri_mCherry/_cyto_GFP *E. coli* MG1655)^18^. In addition, we included a naturally impermeable DNA dye (Sytox) to measure inner membrane destabilization. We treated bacteria with 1% serum or Δlysozyme serum and measured the fluorescence intensity. A reduced mCherry signal, an increase in Sytox intensity and a reduced viability was observed in the presence *and* in the absence of lysozyme, showing that lysozyme is not essential to damage both bacterial membranes or kill *E. coli* in serum (**Fig. 1c** and **SFig. 1b**). Instead, outer and inner membrane damage caused by serum were dependent on the MAC, since both membranes remained intact and bacteria survived in the presence of OmCI (**Fig. 1c** and **SFig. 1b**). Altogether, these data show that although MAC-dependent damage to both bacterial membranes does not depend on lysozyme, lysozyme is essential to further degrade the bacterial cell wall in serum causing changes in FSC/SSC measured by flow cytometry.

### The MAC and lysozyme in serum trigger alterations in the morphology of E. coli

Next, we questioned whether the bacteria that were not detectable anymore using our flow cytometry settings in figure 1 were completely disintegrated, or whether their shape changed in such a way that they appeared differently in the FSC/SSC plots. To track the changes in FSC/SSC of serum-exposed bacteria, we labeled their LPS with a DBCO-Cy3 dye through click-chemistry with a metabolically incorporated KDO-azide (**Fig. 2a**)^24^. In addition, we removed the SSC threshold in our flow cytometry settings, to be able to detect smaller particles in the samples. Cy3-labeled bacteria were exposed to serum or Δlysozyme serum with or without purified lysozyme, after which the number of Cy3-positive particles was quantified within a fixed volume. In contrast to the particle loss we observed in figure 1, the total number of Cy3-positive events remained similar in all conditions (**Fig. 2b**). This suggests that the total number of bacterial particles detected by the flow cytometer is not altered by incubation with serum. When we further analyzed the FSC/SSC of these Cy3-positive particles, we again observed that bacteria exposed to serum have an altered size and shape, as evidenced by a changed FSC/SSC (**Fig. 2c**), which was less pronounced in the absence of lysozyme. Instead, the number of Cy3-positive particles inside the FSC/SSC gate drastically decreased when extra lysozyme was added to the Δlysozyme serum (**Fig. 2c, d**). Altogether, these data show that the combination of the MAC and lysozyme in serum triggers alterations in the shape and size of *E. coli*, which we observe by a FSC/SSC shift compared to non-treated bacteria. Although these bacteria are not completely disintegrated, the cell wall is damaged in such a way that it drastically changes the bacterial morphology.

**Figure 2:**
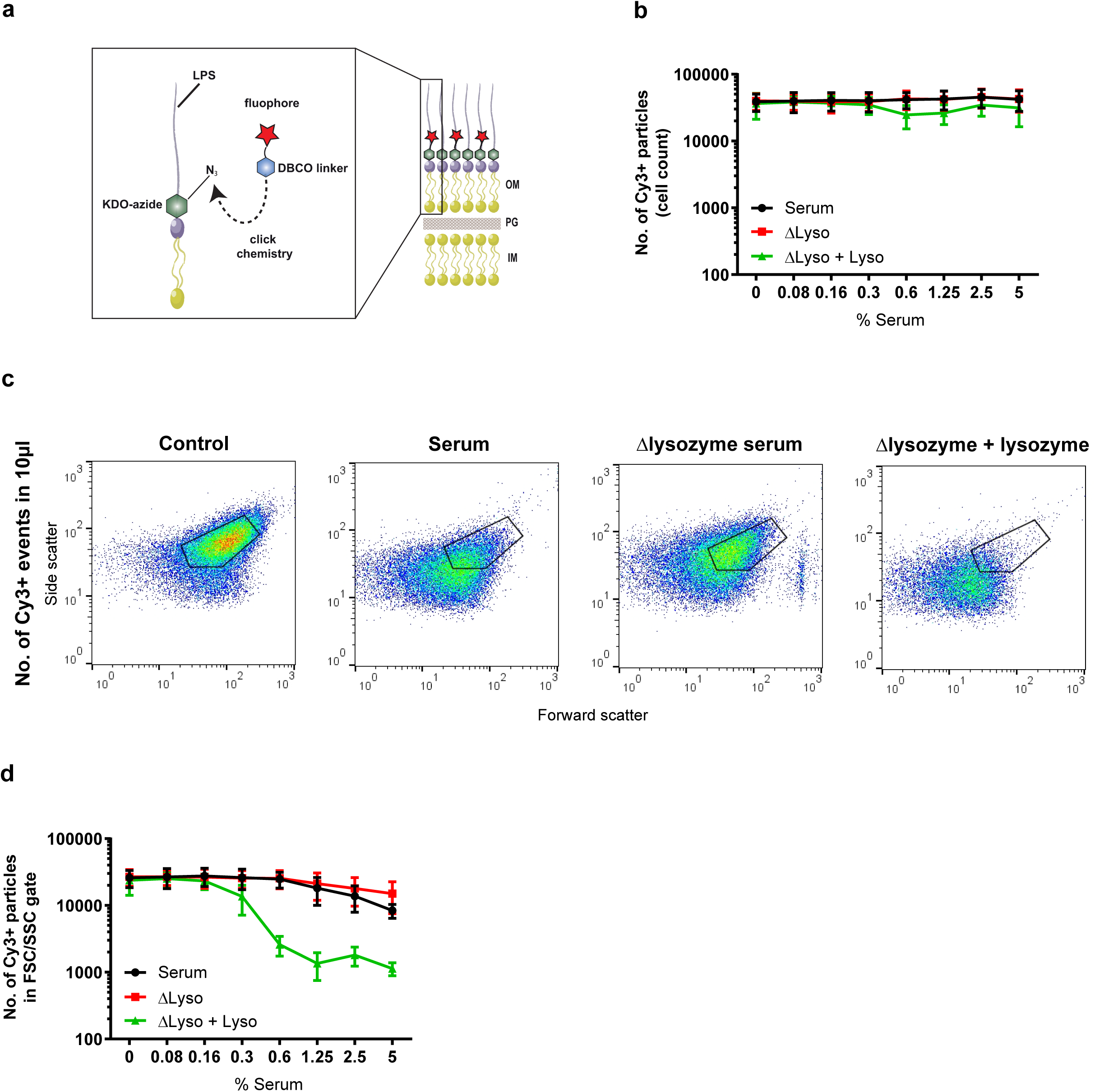
The MAC and lysozyme in serum trigger alterations in the morphology of *E. coli*. **A**) Schematic representation of Cy3-labeling of *E. coli* LPS via click chemistry. Bacteria are incubated with KDO-azide (green), which is incorporated into LPS. A DBCO (blue) linked to the fluorophore Cy3 (red) can subsequently react with the azide group via click chemistry. OM = outer membrane, PG = peptidoglycan, IM = inner membrane. **B**) The number of Cy3-positive *E. coli* cells in 10µl sample after exposure to a concentration range of serum or Δlysozyme serum with or without 5 µg/ml lysozyme. Samples were measured without a SSC threshold. **C**) Flow cytometry plots (FSC/SSC) of Cy3-positive *E. coli* particles in 10 µl sample after exposure to buffer, 5% nhs, or 5% Δlysozyme serum with or without 5 µg/ml lysozyme for 60 minutes at 37°C. **D**) Cy3-labeled *E. coli* was treated with a concentration range of serum or Δlysozyme serum with or without 5 µg/ml lysozyme for 60 min 37°C. For each condition, the number of Cy3-positive particles within the FSC/SSC gate of untreated bacteria was quantified in 10µl. **B, D**) Data represent mean ±SD of 3 independent experiments. **C**) Flow cytometry plots represent data of three independent experiments.

### The MAC and lysozyme alter the cell shape of E. coli from rod-shaped to spherical

To further validate our observations by flow cytometry, we then aimed to visualize the effect of serum on the morphology of *E. coli* in more detail by confocal microscopy. To do so, we treated _peri_mCherry/_cyto_GFP *E. coli* with serum and analyzed the morphology in time. We added an impermeable DNA dye (ToPro-3) to monitor inner membrane destabilization. The inner membrane of these bacteria was efficiently damaged after approximately 60 minutes of exposure to serum at room temperature (RT). After 90 minutes of serum exposure, the cell shape of almost all bacteria had changed from rod-shaped to spherical, suggesting that also the peptidoglycan layer was affected (**Fig. 3a**). We subsequently addressed the role of lysozyme in these experiments by exposing bacteria to Δlysozyme serum with and without purified lysozyme. When bacteria were exposed to Δlysozyme serum that was supplemented with purified lysozyme, bacteria rapidly lost their rod-shape, followed by a loss of GFP and Topro-3 positive particles (**Fig. 3b**). In line with observations in figure 1 and 2, this happened more efficient than in normal serum. In contrast, although we observed efficient inner membrane damage in Δlysozyme serum within 40 minutes, these bacteria remained rod-shaped within the measured time-frame (**Fig. 3b**). Lysozyme-dependent alterations in cell shape were also dependent on the presence of MAC pores, since no shape loss was observed in the presence of the C5 inhibitors OmCI and Eculizumab (**Fig. 3c**). To better visualize the differences in cell shape of these bacteria, 3D reconstructions were generated of the conditions described in figure 3b and c (**SFig. 2**). Altogether, these confocal images confirm that the MAC in serum allows lysozyme to trigger alterations in the shape of *E. coli* from rod-shaped to spherical.

**Figure 3:**
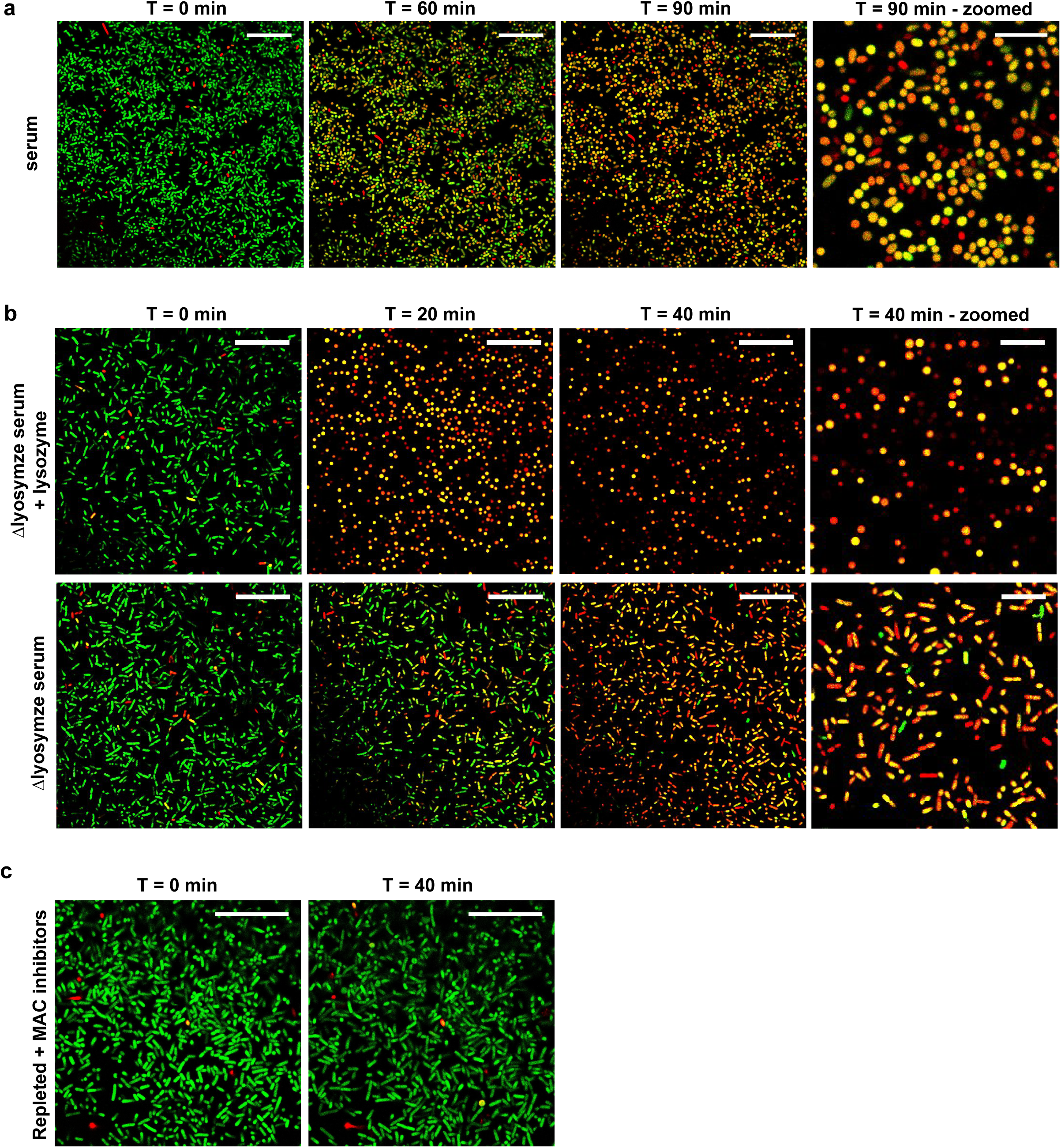
The MAC and lysozyme alter the cell shape of *E. coli* from rod-shaped to spherical. Confocal microscopy images of _Peri_mCherry/_cyto_GFP *E. coli* bacteria that were immobilized onto poly-L-lysin coated coverslips and treated with **A**) 5% serum or **B**) 5% Δlysozyme serum with (top) or without (bottom) 5 µg/ml lysozyme. All incubations are in the presence of To-pro-3 as a readout for inner membrane damage. Images were taken **A**) after 0, 60 and 90 minutes or **B**) after 0, 20 and 40 minutes at room temperature. In **C**), bacteria were treated with 5% Δlysozyme serum with 5 µg/ml lysozyme in the presence of 20 µg/ml OmCI and eculizumab to block MAC formation. Scale bars of non-zoomed images: 25 µm, Scale bars of zoomed images: 10 µm. Images represent data of at least two independent experiments.

### A completely assembled MAC pore is required for cell wall degradation by lysozyme

Next, we wanted to exclude the contribution of other serum components to cell wall degradation in serum and aimed to test whether a complete MAC pore is required for lysozyme-dependent cell wall destruction. To do so, we pre-labeled bacteria with C9-depleted serum (ΔC9 serum), leading to C5b-8 complex formation on the bacterial surface. Since no mCherry (26.7 kDa protein) leaks out through C5b-8 pores^18^, we assume that no serum proteins can enter the periplasmic space at this stage. C5b-8-labeled bacteria were washed, after which purified C9 was added in the presence or absence of lysozyme. A decrease in the number of particles inside the FSC/SSC gate was observed when lysozyme was added in combination with C9 (**Fig. 4a**). In contrast, although the bacterial inner membrane was efficiently damaged in the presence of C9 only, no alteration in FSC/SSC were detected (**Fig. 4a, b**). Lysozyme by itself did not trigger inner membrane damage or alterations in the FSC/SSC on bacteria with C5b-8 complexes (**Fig. 4a, b**). Next, we added a titration of C9 to bacteria with C5b-8 complexes in the presence or absence of lysozyme, and monitored mCherry leakage and particle count. Lysozyme-induced particle loss coincided with the leakage of mCherry from the periplasm (**Fig. 4c**). The particle count inside the FSC/SSC gate remained stable with increasing concentrations of C9 in the absence of lysozyme (**Fig. 4c**), confirming that the MAC does not trigger a change in morphology by itself^18^. Taken together, these data indicate that completely assembled MAC pores allow proteins such as mCherry and lysozyme to cross the bacterial outer membrane.

**Figure 4:**
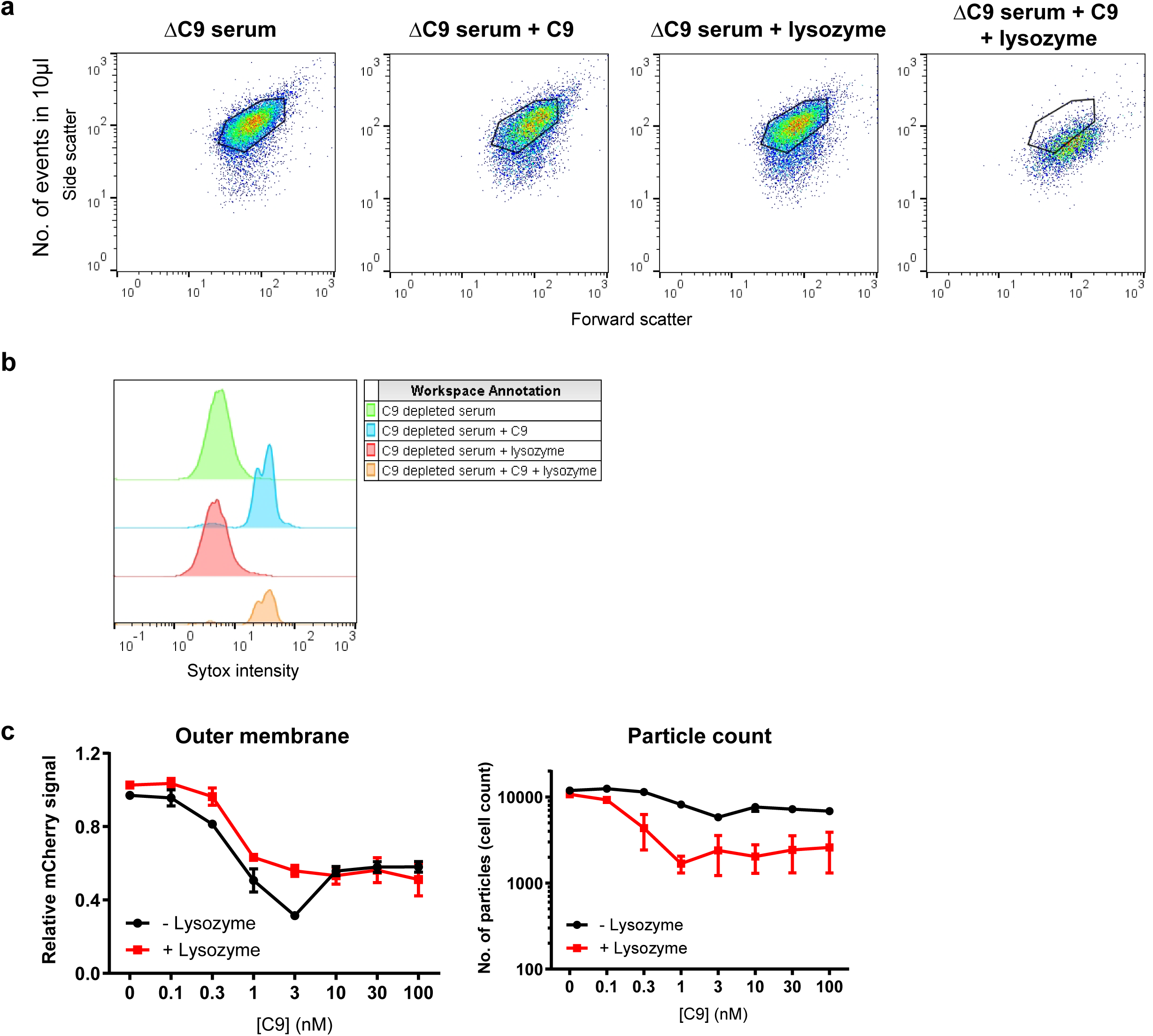
A completely assembled MAC pore is required for cell wall degradation by lysozyme. **A**) Flow cytometry plots (FSC/SSC) and **B**) inner membrane damage (Sytox blue intensity) of _Peri_mCherry/_cyto_GFP *E. coli* that were pre-treated with ΔC9 serum and, after washing, exposed to 100 nM C9 in the presence or absence of 5 µg/ml lysozyme. **C**) Outer membrane damage (left: relative mCherry signal) and particle count (right) of _Peri_mCherry/_cyto_GFP *E. coli* that were treated with ΔC9 serum, and after washing exposed to a concentration range of C9 in the presence or absence of 5 µg/ml lysozyme. Particle count represents the number of particles in the FSC/SSC gate of untreated bacteria. **A, B**) Flow cytometry plots and histograms represent data of three independent experiments. **C**) Data represent mean ±SD of 3 independent experiments.

### The MAC sensitizes E. coli to antimicrobial proteins with different effector functions

As extensively described above, outer membrane permeabilization by the MAC sensitizes *E. coli* to cell wall degradation by lysozyme. Next, we questioned whether this is also true for other antimicrobial proteins, with a completely different function. Therefore, we included an antimicrobial protein that hydrolyzes the cytoplasmic membrane of bacteria, Type IIA secreted phospholipase A2 (hGIIA)^21^. Similar to lysozyme, hGIIA is considered specific for Gram-positive bacteria^25^. To this end, bacteria were pre-labeled bacteria with ΔC9 serum and exposed to C9, in the presence or absence of lysozyme or recombinant hGIIA. As described above, C9 alone did not alter the shape of these bacteria, whereas a change in FSC/SSC was observed when a combination of C9 and lysozyme was added (**Fig. 5a, b**). Although some particles were lost from the FSC/SSC gate with only hGIIA, this was much more efficient when a combination of hGIIA and C9 was added (**Fig. 5b**). Bacteria that were still detected after treatment with a combination of hGIIA and C9 appeared within the FSC/SSC of untreated bacteria (**Fig. 5a**). Almost all events were lost when bacteria were treated with a combination of C9, lysozyme and hGIIA (**Fig. 5a, b**). We then tested the effect of the MAC, hGIIA, lysozyme or a combination of those on the level of cytoplasmic GFP of the (remaining) bacteria. Although hGIIA alone slightly decreased the intensity of cytoplasmic GFP, the addition of increasing concentrations of C9 drastically decreased the GFP signal of these bacteria. In contrast, whereas lysozyme triggered efficient shape loss in the presence of C9, these bacteria remained GFP positive. In line with previous observations, the MAC alone also had no effect on the level of cytoplasmic GFP (**Fig. 5c**)^10^. Altogether, we conclude that the MAC sensitizes *E. coli* to antimicrobial proteins with different effector functions, acting on the peptidoglycan layer (lysozyme) and on the bacterial inner membrane (hGIIA).

**Figure 5:**
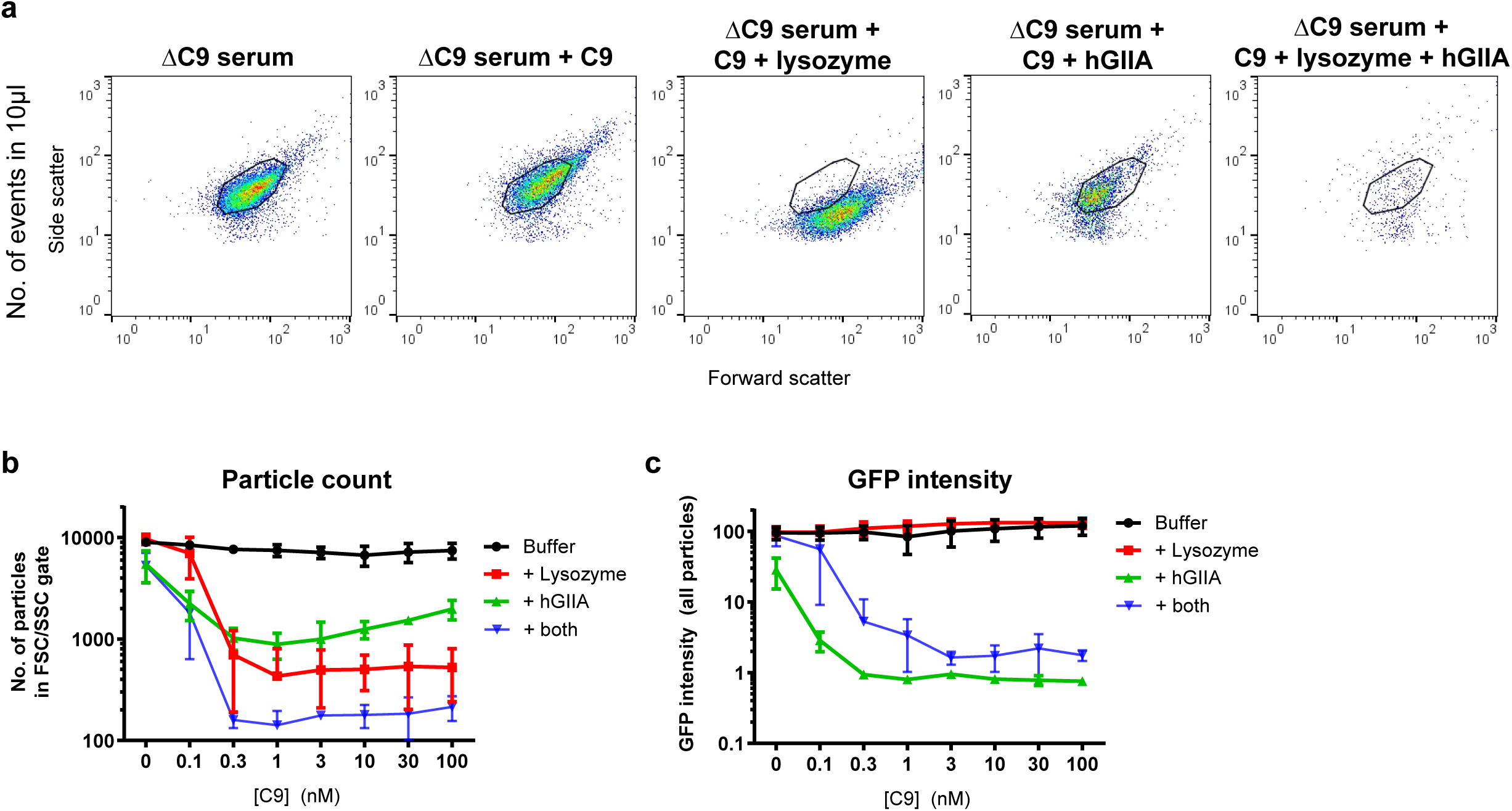
The MAC sensitizes *E. coli* to antimicrobial proteins with different effector functions. **A**) Flow cytometry plots (FSC/SSC), **B**) particle count and **C**) GFP intensity of _Peri_mCherry/_cyto_GFP *E. coli* that was pre-treated with ΔC9 serum and, after washing, exposed to a concentration range of C9 in the presence or absence of 5 µg/ml lysozyme and/or 1µg/ml hGIIA. In **A**, the flow cytometry plots of buffer or 100 nM C9 are depicted. **B**) A gate was set on untreated bacteria as depicted in **A**, after which the number of particles was counted within those gates. In **C**), the GFP intensity (Geomean) of the total number of events is presented. All incubations were performed for 30 minutes at 37°C. **A**) Flow cytometry plots represent data of three independent experiments. **B, C**) Data represent mean ±SD of 3 independent experiments.

### The MAC sensitizes E. coli to killing by neutrophils

For intracellular killing by neutrophils, Gram-negative bacteria are taken up into phagolysosomes that contain a large number of antimicrobial proteins and peptides^16,26^. Several of these proteins, including lysozyme and hGIIA, are considered inactive against Gram-negative bacteria and require other factors in the phagolysosome to first damage the bacterial outer membrane^12,27–30^. Given that the MAC efficiently sensitizes *E. coli* to antimicrobial proteins in serum, we hypothesize that it may also sensitize bacteria to antimicrobial factors inside neutrophils. To test this, we treated _Peri_mCherry/_cyto_GFP *E.coli* with Δlysozyme serum to allow MAC formation while maintaining the cell shape (**Fig. 6a**), and exposed them to neutrophils to allow phagocytosis. Intracellular bacteria were efficient degraded, as evidenced by the diffused GFP signal and the absence of clear rod shape bacteria (**Fig. 6a**). In serum, bacteria are efficiently labeled with C3b molecules, which enhances phagocytosis and thereby facilitates bacterial killing by neutrophils^31,32^. However, the contribution of MAC pores to killing by neutrophils is less well understood. To directly study the effect of MAC pores on sensitizing bacteria to neutrophils, we incubated _Peri_mCherry/_cyto_GFP *E.coli* with ΔC5 serum to label the bacterial surface with C5 convertase as previously described^18^. These bacteria were then exposed to buffer, or to purified C5-C9 to allow pore formation. When convertase-labeled bacteria were internalized by neutrophils, bacteria remained rod-shaped within the measured time-frame (**Fig. 6b**). Instead, when convertase-labeled bacteria were first exposed to purified MAC components (C5-C9) and then internalized by neutrophils, bacteria lost their rod shape (**Fig. 6b**). Importantly, a higher laser intensity was needed to detect the GFP signal in the samples with MAC components. Besides the fact that GFP is more diffuse in these conditions, it might also be degraded inside the phagolysosome. To test whether a full MAC pore is required to sensitize bacteria to neutrophils, we included a condition in which C9 was omitted, to allow MAC formation up to C8. When these bacteria were phagocytosed, they still appeared as rod-shaped particles inside the neutrophil (**Fig. 6b**), showing that a complete MAC pore (C5b-9) is required to sensitize *E. coli* to intracellular degradation by neutrophils.

**Figure 6:**
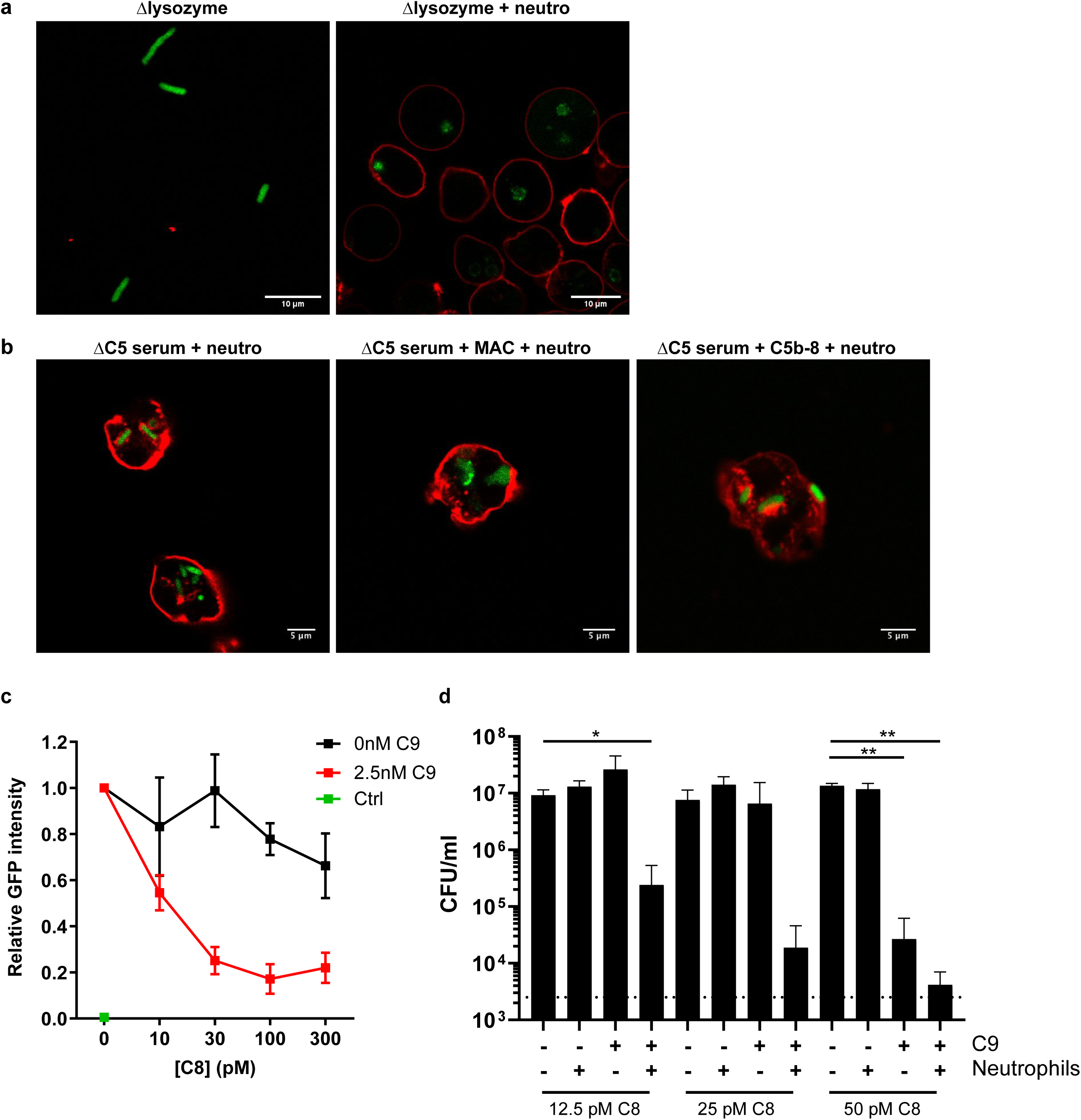
The MAC sensitizes *E. coli* to killing by neutrophils. **A**) Confocal microscopy images of _Peri_mCherry/_cyto_GFP *E. coli* (green) treated with 5% Δlysozyme serum for 60 minutes at 37°C. After washing, bacteria were incubated with buffer (left) or neutrophils (red, right) for 20 minutes at 37°C, fixed in 1.5% paraformaldehyde and imaged. **B**) Confocal images of _Peri_mCherry/_cyto_GFP *E. coli* (green) that were pre-labeled with 10% ΔC5 serum (for deposition of C5 convertases), washed and exposed to buffer, C5-C8 or C5-C9 for 30 minutes at 37°C. 10 nM C5 and C6, 20 nM C7 and C8 and 100 nM C9 was used. After washing, bacteria were exposed to neutrophils for 20 minutes at 37°C, fixed in 1.5% paraformaldehyde and imaged. Images were obtained in separate experiments in which the 488-laser settings were adjusted to the GFP intensity. **A, B**) Neutrophils membranes (red) were stained with Alexa-647 labeled Wheat Germ Agglutinin. **C**) Phagocytosis (relative GFP intensity) of _Peri_mCherry/_cyto_GFP *E. coli* by neutrophils. Bacteria were pre-labeled with 10% ΔC8 serum and, after washing, exposed to a concentration range of C8 in the presence or absence of 2.5 nM C9 for 30 minutes at 37°C. After washing bacteria were incubated with neutrophils for 20 minutes at 37°C, after which the GFP signal of the neutrophils was analyzed by flow cytometry. GFP intensity was normalized against bacteria that were treated with ΔC8 serum only. Neutrophils without bacteria (green dot) served as basal value for the GFP signal. **D**) Bacterial viability (CFU/ml) of *E. coli* that was pre-treated with MAC components similar to **C**, and exposed to buffer or neutrophils. After 20 minutes, neutrophils were lysed for 15 minutes in MQ, after which bacterial survival was determined (CFU/ml). **C, D**) Bacteria/neutrophil ratio used: 10/1. **D**) Statistical analysis was done between samples within each C8 concentration using a one-way ANOVA with Geisser-Greenhouse correction and a Tukey’s multiple comparisons test and displayed only when significant as *P ≤ 0.05 or **P ≤ 0.01.

We next performed flow cytometry experiments to study the role of the MAC inside neutrophils in a more high-throughput way. We carefully titrated the number of MAC pores on _Peri_mCherry/_cyto_GFP bacteria by exposing them to ΔC8 serum and, after washing, a concentration range of C8 in the presence or absence of C9. These bacteria were subsequently exposed to neutrophils. In each experiment, the GFP intensity of neutrophils exposed to bacteria treated with ΔC8 serum was used as a reference value (**Fig. 6c**). The GFP signal within the neutrophil population drastically decreased when pre-labeled bacteria were incubated with both C8 and C9. In contrast, it remained relatively stable when only C8 was added (**Fig. 6c**). This correlates with the confocal experiments, in which a higher 488-laser intensity was required to detect GFP inside neutrophils when bacteria were treated with full MAC pores. We next repeated the experiment with bacteria of which the LPS was labeled with Cy3 (as described above) to verify that the efficiency of phagocytosis was unaffected by the pre-treatment with different combinations of MAC components. The Cy3 signal of the neutrophil population was equal in each condition, suggesting that the same amount of bacteria was phagocytosed and that Cy3 is less sensitive to losing its fluorescence inside neutrophils than GFP (**SFig. 3**). Finally, we used the same experimental setup to assess whether the MAC also increases bacterial killing (CFU/ml) by neutrophils. When only C8 was added to bacteria pre-treated with ΔC8 serum, no killing was observed in the presence or absence of neutrophils (**Fig. 6d**). At the highest concentration of C8 (50 pM), *E. coli* was efficiently killed in the presence of C9, and therefore only a slight additive effect of neutrophils could be measured (**Fig. 6d**). However, at the C8 concentrations where addition of C9 or neutrophils alone was not sufficient to kill these bacteria (12.5 and 25 pM), the combination of both triggered efficient killing of these bacteria (**Fig. 6d**). Altogether, these results show that the MAC enhances cell wall degradation and killing of *E. coli* inside neutrophils.

## Discussion

Infections with Gram-negative bacteria form an increasing problem for human health, which can partly be attributed to the presence of an outer membrane that is selectively permeable to antibiotics and endogenous antimicrobials. Therefore, combination therapy of antibiotics and outer membrane permeabilizing agents have become more attractive over the last decades^5–9^. Additionally, our own immune system can damage the bacterial outer membrane in such a way that it sensitizes bacteria to naturally ineffective antibiotics. Specifically, MAC pores efficiently disrupt the bacterial outer membrane, allowing antibiotics to enter the periplasmic space and fulfill their antimicrobial functions^10^. Our recently developed fluorescent reporter system enabled us to unravel in detail how the complement system kills Gram-negative bacteria. These tools are essential to discriminate between different types of membrane damage and to study how these influence bacterial viability and cell wall integrity. By labeling different cell compartments and the bacterial cell membrane (LPS), we are now able to study membrane damage and shape changes by flow cytometry and confocal microscopy. Using these tools, we previously demonstrated that the MAC forms pores in the bacterial outer membrane that trigger destabilization of the bacterial inner membrane, which is essential for bacterial killing. However, we noticed that the complement system alone does not affect the cell morphology of Gram-negative bacteria^18^, suggesting that other factors are required for further cell wall degradation. Here we show that the MAC plays a crucial role in sensitizing Gram-negative bacteria to other human immune factors, such as antimicrobial proteins in- and outside phagocytes.

One of these factors is lysozyme, an antimicrobial protein that degrades peptidoglycan. Using flow cytometry, we observed that the MAC and lysozyme in serum are both required to trigger alterations in the morphology of *E. coli.* These findings were verified by confocal microscopy, in which we observed that the MAC and lysozyme together affect the cell wall in such a way that *E. coli* bacteria change from rods to spheres. These findings correlate with previous electron microscopy studies, showing that the MAC can kill bacteria, but that a combination of the MAC and lysozyme damages the bacterial cell wall more severely^33–36^. The peptidoglycan layer of Gram-negative bacteria is essential for the integrity of the bacterial cell wall and to withstand turgor pressure from the cytoplasmic space^37^. Degradation of this layer by lysozyme could therefore explain why these bacteria lose their rod-shaped morphology. Although lysozyme is generally known as an antimicrobial protein against Gram-positive bacteria, several Gram-negative species (among which *E. coli* and *N. gonorrhoeae*) have evolved resistance mechanisms against these enzymes^38–40^. The fact that lysozyme plays a crucial role in cell wall degradation of Gram-negative bacteria in the presence of the complement system may explain why some of these bacteria have developed evasion strategies.

The MAC also sensitizes *E. coli* to hGIIA, which hydrolyses phospholipid membranes^21^. A combination of MAC pores and hGIIA triggered efficient leakage of cytoplasmic proteins, and a loss of particles detected by flow cytometry. The fact that MAC pores in the outer membrane allow both lysozyme and hGIIA to severely damage the bacterial cell envelope suggests that this principle applies to a broad range of antimicrobial proteins that are normally inactive against Gram-negative bacteria. Lysozyme and hGIIA are both secreted by epithelial cells and by immune cells that are recruited towards the site of infection^13,21^. The concentration of lysozyme in normal serum is approximately 1 µg/ml^18^, and increases in body fluids under inflammatory conditions^41^. The normal serum concentration of hGIIA is approximately 4 ng/ml^42^ and rises up to 1µg/ml in inflamed serum^43^. Although the concentrations of these proteins have been measured in body fluids in “inflamed conditions”, it remains challenging to determine what concentrations would be relevant to mimic the conditions at the site of infection or inside immune cells. These will most probably vary between individuals, depend on the location of the infection and on the immune status of the patient. To the best of our knowledge, there are no studies available that directly compare lysozyme or hGIIA concentrations in serum of healthy donors and serum of patients that have an infection with a Gram-negative bacterium. The protein concentrations we used in this study should therefore be considered as a proof of principle.

At the site of infection, bacteria can also be internalized by immune cells. Phagocytosed bacteria are subsequently exposed to high concentrations of intracellular lysozyme, hGIIA and other antimicrobial peptides and proteins. Among these proteins are lactoferrin, defensins and bactericidal permeability increasing protein (BPI), which can all enhance outer membrane permeability^12,27–30^. The complement system enhances phagocytosis of Gram-negative bacteria by depositing C3b on the surface, which interacts with complement receptors on the neutrophil surface^31,32^. However, within the time-frame of our experiments, internalized C3b-labeled bacteria were not degraded or killed by neutrophils, suggesting that this process is relatively inefficient. In contrast, internalized bacteria were efficiently degraded and killed when these were pre-treated with full MAC pores (C5b-9) that allow intracellular proteins to cross the outer membrane^18^. These results suggest that the complement system does not only enhance phagocytosis via the deposition of C3b molecules, but that the MAC also plays a crucial role in degradation of Gram-negative bacteria by neutrophils. Vice versa, intracellular or secreted antimicrobial proteins that enhance the outer membrane permeability of bacteria (such as BPI) may also enhance the efficiency with which MAC pores insert into the bacterial outer membrane and kill bacteria^9^. Although the tested *E. coli* strain is sensitive to killing by certain concentrations of MAC pores alone, we showed that these bacteria are also efficiently killed inside neutrophils when opsonized with non-bactericidal concentrations of MACs. It is likely that several factors in the phagolysosome, among which lysozyme and hGIIA^19^ are involved in the killing of MAC-opsonized bacteria. However, since killing of *E. coli* by neutrophils is only enhanced in the presence of all MAC components, it likely depends on proteins that need to cross the outer membrane.

We hypothesize that MAC-dependent cell wall degradation of bacteria by lysozyme, hGIIA and other antimicrobial proteins may facilitate and accelerate clearance of these particles from the body, for example by neutrophils. Since the shape of internalized particles influences immune cell activation^44^, this may also play a role in antigen presentation and stimulation of a proper adaptive immunity response. Faster clearance of Gram-negative bacteria from the body could prevent ongoing immune activation on the surface of these bacteria that are already killed. However, further research is needed to show the exact relevance of each step of cell envelope degradation on immune-mediated clearance of bacteria.

The fact that MAC-dependent outer membrane damage sensitizes Gram-negative bacteria to cell envelope degradation by lysozyme and hGIIA suggests that also other outer membrane permeabilizing agents can synergize with human immune factors. In this study, we showed that the MAC sensitizes *E. coli* to the antimicrobial actions of lysozyme, hGIIA and neutrophils. However, we propose that this mechanism applies to a broad range of antimicrobial proteins, peptides and antibiotics that normally fail to pass the outer membrane of Gram-negative bacteria^10,45^. The synergy between MAC pores and antimicrobial proteins might even be more crucial to kill bacterial strains on which MAC pores form in the outer membrane that fail to damage the inner membrane and kill these bacteria.

## Material and methods

### Serum and reagents

Pooled normal human serum (NHS) was obtained from healthy volunteers as previously described^46^. Human neutrophils were isolated on the day of the experiment from freshly drawn heparinized blood of healthy volunteers using density gradient centrifugation^47^. Complement factor C8 and sera depleted of C5, C8 or C9 were obtained from Complement Technology. His-tagged C5, C6, C7 and C9 and OmCI were expressed in and purified from HEK293E cells (U-Protein Express)^23^. Lysozyme-depleted serum was prepared by affinity depletion using an LprI column and checked for complement activity, as described in^18^. Human neutrophil lysozyme was obtained from RayBiotech. To obtain _Peri_mCherry/_cyto_GFP *E. coli*, MG1655 was transformed with a pPerimCh plasmid, containing a constitutively expressed periplasmic mCherry and a L-arabinose inducible cytosolic GFP^18^. GFP expression was induced using 0.1% L-arabinose. Recombinant hGIIA was produced as previously described^48^. Eculizumab was kindly provided by Frank Beurskens (Genmab, Utrecht, The Netherlands). KDO-azide was synthesized according to a reported procedure^49^.

### LPS labeling via click-chemistry

A single colony of *E. coli* MG1655 was grown to stationary phase in Lysogeny Broth (LB) medium. Bacteria were subcultured by diluting 1/100 in LB medium in the presence of 2 mM KDO-azide and incubated overnight (o/n), shaking at 37°C. The next day, bacteria were washed three times in RPMI 1640 medium (Thermofisher) containing 0.05% human serum albumin (RPMI-HSA) and incubated with 50 µM DBCO-Cy3 for 2.5 hours, shaking at 4°C. Bacteria were washed three times in RPMI-HSA and resuspended in RPMI-HSA at OD_600_∼0.1.

### Flow cytometry and bacterial viability assay

In all experiments, *E. coli* was grown o/n in LB medium. _Peri_mCherry/_Cyto_GFP *E. coli* were grown in the presence of 100 µg/ml ampicillin. The next day, subcultures were grown to mid-log phase (OD_600_∼0.5) in the presence of 0.1% L-arabinose, washed and resuspended in RPMI-HSA. Bacteria (50 µl) of OD_600_∼0.025 were mixed with buffer, NHS or Δlysozyme serum with or without 5 µg/ml lysozyme. For assays using purified complement components, bacteria were pre-treated with 10% ΔC5, ΔC8 or ΔC9 serum for 30 minutes at 37°C, washed three times and further incubated with buffer or the remaining MAC components for 30 or 60 minutes, as indicated in the figure legend. Also the concentrations of purified MAC components are specified in the figure legends. When indicated, 5 µg/ml lysozyme and 1 µg/ml hGIIA was added to these incubations. One µM Sytox blue or Sytox Green (ThermoFisher) was used to measure inner membrane damage. The number of particles, or the GFP, Sytox blue or green, mCherry or Cy3 intensity was measured by a MACSQuant flow cytometer (Miltenyi Biotech). For this, either a fixed volume (10 µl) or a fixed number of particles (10.000) was measured, with or without a SSC threshold, as indicated in the figure legends. To determine bacterial viability, samples were serially diluted into PBS and plated onto LB agar plates. Colonies were counted after overnight incubation.

### Neutrophil phagocytosis and killing assay

Non-labeled or Cy3-labeled _Peri_mCherry/_Cyto_GFP bacteria were treated as described in the figure legends, and incubated with neutrophils at a bacteria/neutrophil ratio of 10:1 to allow phagocytosis for 20 minutes shaking at 37°C. For neutrophil phagocytosis assays, samples were diluted twenty times in RPMI + 0.05% HSA after which the GFP or Cy3 signal of a fixed number of neutrophils (10^5^) was analyzed by flow cytometry (MACSQuant, Miltenyi Biotech). To determine bacterial viability, samples were diluted 1:20 in MilliQ and incubated for 15 minutes to allow neutrophil lysis. Afterwards, samples were serially diluted into PBS and plated onto LB agar plates. Colonies were counted after overnight incubation.

### Confocal microscopy

For the time-course experiments, _Peri_mCherry/_Cyto_GFP *E. coli* were grown in the presence of L-arabinose as described above, washed three times in PBS and concentrated to OD_600_∼2 in PBS. Bacteria were immobilized on a poly-L-lysine (0.01%, Sigma-Aldrich) covered 8 well µ-slide chamber (Ibidi) for 45 minutes. Chambers were rinsed three times with PBS after which RPMI-HSA containing 1 µM To-pro-3 (Thermofisher) was added. A T=0 image was taken, after which 5% normal serum or Δlysozyme serum with or without 5 µg/ml purified lysozyme was added. The Δlysozyme serum + lysozyme was imaged with and without 20 µg/ml OmCI and Eculizumab. GFP and To-pro-3 intensity was measured after the indicated time-pointes at room temperature. To image GFP bacteria inside neutrophils, the samples were prepared as described above and fixed in 1.5% paraformaldehyde. Neutrophil membranes were stained for 15 minutes with 2 µg/ml Alexa Fluor 647-conjugated Wheat Germ Agglutinin (ThermoFisher). Samples were concentrated and dried onto 1% agar pads. All images were obtained using a Leica SP5 confocal microscope with a HCX PL APO CS 63×/1.40–0.60 OIL objective (Leica Microsystems, the Netherlands). GFP was measured using the 488 laser, Alexa-647 and To-pro-3 were imaged using the 647 laser. Both lasers were used in combination with the appropriate emission filter settings.

### Data analysis and statistics

Flow cytometry data was analyzed using FlowJo (version 10). Graphpad 6.0 was used for graph design and statistical analysis. Statistical analysis was done using a one-way ANOVA with the appropriate corrections for multiple comparisons as indicated in the figure legends. Three experimental replicates were performed to allow statistical testing. Confocal images were processed in Fiji.

## Supporting information

Supplementary information - Heesterbeek et al

## Acknowledgements

We would like to acknowledge Kok van Kessel for proofreading the manuscript and for useful discussions. This work was funded by: an ERC Starting grant (639209-ComBact, to S.H.M.R) and the Molecular immunology HUB (eSTIMATE). This work was supported by Vidi grants from the Dutch Research Council (NWO) to N.M.v.S. (917.13.303) and T.W. (723.014.005).

## References

1. Tacconelli, E. et al. Discovery, research, and development of new antibiotics: the WHO priority list of antibiotic-resistant bacteria and tuberculosis. Lancet. Infect. Dis. 18, 318–327 (2018).

2. Silhavy, T. J., Kahne, D. & Walker, S. The bacterial cell envelope. Cold Spring Harb. Perspect. Biol. 2, a000414 (2010).

3. Cabeen, M. T. & Jacobs-Wagner, C. Bacterial cell shape. Nat. Rev. Microbiol. 3, 601–610 (2005).

4. Delcour, A. H. Outer Membrane Permeability and Antibiotic Resistance. Biochim Biophys Acta. 1794, 808–816 (2009).

5. Stokes, J. M. et al. Pentamidine sensitizes Gram-negative pathogens to antibiotics and overcomes acquired colistin resistance. Nat. Microbiol. 2, 17028 (2017).

6. MacNair, C. R. et al. Overcoming mcr-1 mediated colistin resistance with colistin in combination with other antibiotics. Nat. Commun. 9, 458 (2018).

7. Bolla, J., Alibert-franco, S., Handzlik, J. & Chevalier, J. Strategies for bypassing the membrane barrier in multidrug resistant Gram-negative bacteria. FEBS Lett. 585, 1682–1690 (2011).

8. Jammal, J., Zaknoon, F., Kaneti, G., Goldberg, K. & Mor, A. Sensitization of Gram-negative bacteria to rifampin and OAK combinations. Sci. Rep. 5, 9216 (2015).

9. Vaara, M. & Vaara, T. Sensitization of Gram-negative bacteria to antibiotics and complement by a nontoxic oligopeptide. Nature vol. 303 526–528 (1983).

10. Heesterbeek, D. A. C. et al. Complement-dependent outer membrane perturbation sensitizes Gram-negative bacteria to Gram-positive specific antibiotics. Sci. Rep. 9, 3074 (2019).

11. Segal, A. W. How neutrophils kill microbes. Annu. Rev. Immunol. 23, 197–223 (2005).

12. Nathan, C. Neutrophils and immunity: challenges and opportunities. Nat. Rev. Immunol. 6, 173–82 (2006).

13. Ragland, S. A. & Criss, A. K. From bacterial killing to immune modulation: Recent insights into the functions of lysozyme. PLoS Pathog. 13, 1–22 (2017).

14. Weibull, C. The isolation of protoplasts from Bacillus megaterium by controlled treatment with lysozyme. J. Bacteriol. 66, 688–695 (1953).

15. Masschalck, B., Van Houdt, R., Van Haver, E. G. & Michiels, C. W. Inactivation of gram-negative bacteria by lysozyme, denatured lysozyme, and lysozyme-derived peptides under high hydrostatic pressure. Appl. Environ. Microbiol. 67, 339–44 (2001).

16. Urban, C. F., Lourido, S. & Zychlinsky, A. How do microbes evade neutrophil killing? Cell. Microbiol. 8, 1687–96 (2006).

17. Morgan, B. P. et al. CryoEM reveals how the complement membrane attack complex ruptures lipid bilayers. Nat. Commun. 9, 5316 (2018).

18. Heesterbeek, D. A. et al. Bacterial killing by complement requires membrane attack complex formation via surface-bound C5 convertases. EMBO J. 38, e99852 (2019).

19. Madsen, L. M., Inada, M. & Weiss, J. Determinants of activation by complement of group II phospholipase A2 acting against Escherichia coli. Infect. Immun. 64, 2425–2430 (1996).

20. Mannion, B. A., Weiss, J. & Elsbach, P. Separation of sublethal and lethal effects of polymorphonuclear leukocytes on Escherichia coli. J. Clin. Invest. 86, 631–41 (1990).

21. Beers, S. A. et al. The antibacterial properties of secreted phospholipases A2: A major physiological role for the group IIA enzyme that depends on the very high pI of the enzyme to allow penetration of the bacterial cell wall. J. Biol. Chem. 277, 1788–1793 (2002).

22. Vollmer, W. & Seligman, S. J. Architecture of peptidoglycan: more data and more models. Trends Microbiol. 18, 59–66 (2010).

23. Nunn, M. A. et al. Complement Inhibitor of C5 Activation from the Soft Tick Ornithodoros moubata. J. Immunol. 174, 2084–2091 (2005).

24. Dumont, A., Malleron, A., Awwad, M., Dukan, S. & Vauzeilles, B. Click-mediated labeling of bacterial membranes through metabolic modification of the lipopolysaccharide inner core. Angew. Chemie - Int. Ed. 51, 3143–3146 (2012).

25. Weiss, J. P. Molecular determinants of bacterial sensitivity and resistance to mammalian Group IIA phospholipase A2. Biochim. Biophys. Acta 1848, 3072–7 (2015).

26. Borregaard, N. & Cowland, J. B. Granules of the human neutrophilic polymorphonuclear leukocyte. Blood 89, 3503–3521 (1997).

27. Ellison, R. T. & Giehl, T. J. Killing of gram-negative bacteria by lactoferrin and lysozyme. J. Clin. Invest. 88, 1080–1091 (1991).

28. Ganz, T. Defensins: Antimicrobial peptides of innate immunity. Nat. Rev. Immunol. 3, 710–720 (2003).

29. Vaara, M. Agents that increase the permeability of the outer membrane. Microbiol. Rev. 56, 395–411 (1992).

30. Wiese, A., Brandenburg, K., Carroll, S. F., Rietschel, E. T. & Seydel, U. Mechanisms of action of bactericidal/permeability-increasing protein BPI on reconstituted outer membranes of gram-negative bacteria. Biochemistry 36, 10311–9 (1997).

31. Fearon, D. T. Identification of the membrane glycoprotein that is the C3b receptor of the human erythrocyte, polymorphonuclear leukocyte, B lymphocyte, and monocyte. J. Exp. Med. 152, 20–30 (1980).

32. Ross, G. D. & Lambris, J. D. Identification of a C3bi-specific membrane complement receptor that is expressed on lymphocytes, monocytes, neutrophils, and erythrocytes. J. Exp. Med. 155, 96–110 (1982).

33. Donaldson, D. M., Roberts, R. R., Larsen, H. S. & Tew, J. G. Interrelationship between serum beta-lysin, lysozyme, and the antibody-complement system in killing Escherichia coli. Infect. Immun. 10, 657–66 (1974).

34. Feingold, D. S., Goldman, J. N. & Kuritz, H. M. Locus of the action of serum and the role of lysozyme in the serum bactericidal reaction. J. Bacteriol. 96, 2118–2126 (1968).

35. Taylor, P. W. Bactericidal and Bacteriolytic Activity of Serum Against Gram-Negative Bacteria. Microbiol. Rev. 47, 46–83 (1983).

36. Schreiber, R. D., Morrison, D. C., Podack, E. R. & Müller-Eberhard, H. J. Bactericidal activity of the alternative complement pathway generated from 11 isolated plasma proteins. J. Exp. Med. 149, 870–882 (1979).

37. Vollmer, W., Blanot, D. & De Pedro, M. A. Peptidoglycan structure and architecture. FEMS Microbiol. Rev. 32, 149–167 (2008).

38. Ragland, S. A., Humbert, M. V, Christodoulides, M. & Criss, A. K. Neisseria gonorrhoeae employs two protein inhibitors to evade killing by human lysozyme. PLoS Pathog. 14, e1007080 (2018).

39. Liu, Z., García-Díaz, B., Catacchio, B., Chiancone, E. & Vogel, H. J. Protecting Gram-negative bacterial cell envelopes from human lysozyme: Interactions with Ivy inhibitor proteins from Escherichia coli and Pseudomonas aeruginosa. Biochim. Biophys. Acta - Biomembr. 1848, 3032–3046 (2015).

40. Vanderkelen, L. et al. Role of Lysozyme Inhibitors in the Virulence of Avian Pathogenic Escherichia coli. PLoS One 7, 1–8 (2012).

41. Porstmann, B. et al. Measurement of lysozyme in human body fluids: comparison of various enzyme immunoassay techniques and their diagnostic application. Clin. Biochem. 22, 349–55 (1989).

42. Peuravuori, H. J., Funatomi, H. & Nevalainen, T. J. Group I and Group II Phospholipases A2 in Serum in Uraemia. Clin. Chem. Lab. Med. 31, 491–494 (1993).

43. Green, J. A. et al. Circulating phospholipase A2 activity associated with sepsis and septic shock is indistinguishable from that associated with rheumatoid arthritis. Inflammation (1991) doi: 10.1007/BF00917352.

44. Vaine, C. A. et al. Tuning innate immune activation by surface texturing of polymer microparticles: the role of shape in inflammasome activation. J. Immunol. 190, 3525–32 (2013).

45. Wang, G. Human antimicrobial peptides and proteins. Pharmaceuticals 7, 545–594 (2014).

46. Berends, E. T. M. et al. Distinct localization of the complement C5b-9 complex on Gram-positive bacteria. Cell. Microbiol. 15, 1955–1968 (2013).

47. Surewaard, B. G. J., van Strijp, J. A. G. & Nijland, R. Studying interactions of Staphylococcus aureus with neutrophils by flow cytometry and time lapse microscopy. J. Vis. Exp. e50788 (2013) doi: 10.3791/50788.

48. Ghomashchi, F. et al. Preparation of the Full Set of Recombinant Mouse- and Human-Secreted Phospholipases A2. in Methods in Enzymology (2017). doi: 10.1016/bs.mie.2016.10.034.

49. Winzar, R., Philips, J. & Kiefel, M. J. A simple synthesis of C-8 modified 2-keto-3-deoxy-d-manno-octulosonic acid (KDO) derivatives. Synlett (2010) doi: 10.1055/s-0029-1219356.

