## Supplementary information - Heesterbeek et al for "Outer membrane permeabilization by the membrane attack complex sensitizes Gram-negative bacteria to antimicrobial proteins in serum and phagocytes"

**SFigure 1**: **Lysozyme in serum is not essential for bacterial killing, but enhances membrane disintegration in the presence of the MAC. A**) *E. coli* cell count in 10 µl after exposure to a concentration range of serum or Δlysozyme serum with 5 µg/ml lysozyme in the presence or absence of 20 µg/ml OmCI for 60 min 37°C. Flow cytometry settings were similar to Figure 1A and B. **B**) Bacterial viability (CFU/ml) of *E. coli* exposed to buffer, 1% serum with or without 20 µg/ml OmCI or Δlysozyme serum.

**SFigure 2: The MAC and lysozyme alter the cell shape of *E. coli* from rod-shaped to spherical.** 3D reconstructions of confocal microscopy images of $_{Peri}$mCherry/$_{cyto}$GFP *E. coli* bacteria that were immobilized onto poly-L-lysin coated coverslips. Bacteria were treated with Buffer, 5% Δlysozyme serum or 5% Δlysozyme serum with 5 µg/ml lysozyme in the absence or presence of 20 µg/ml OmCI and Eculizumab (Repleted + MAC inhibitors). All incubations are in the presence of To-pro-3 as a readout for inner membrane damage. Images were taken after 45 minutes at room temperature. Scale bars: 10 µm.

**SFigure 3: MAC formation does not influence the number of bacteria phagocytosed by neutrophils.** DBCO-Cy3-labeled *E. coli* were exposed to ΔC8 serum for 30 minutes at 37°C. After washing, bacteria were incubated with buffer or 0.03 nM C8, 2.5 nM C9 or a combination of both. Bacteria were again washed after 30 minutes at 37°C and incubated with neutrophils for 20 minutes at 37°C. Cy3 intensity in neutrophils relative to the buffer control was analyzed by flow cytometry. Statistical analysis was done using a one-way ANOVA with a Dunnett's multiple comparisons test in which each test condition was compared to the buffer control. No significant differences were found.

### SFig 1

a

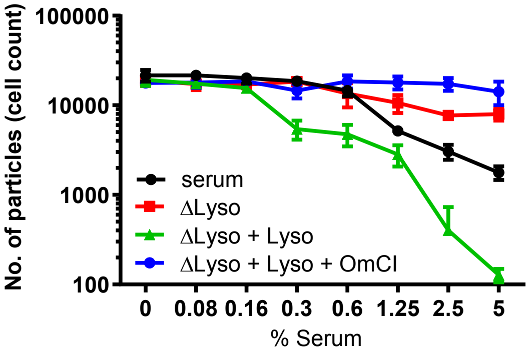
No. of particles (cell count)
100000
10000
1000
100
serum
ΔLyso
ΔLyso + Lyso
ΔLyso + Lyso + OmCI
0
0.08
0.16
0.3
0.6
1.25
2.5
5
% Serum

b

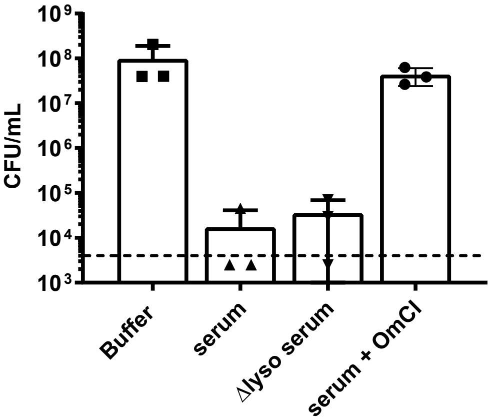
CFU/mL
$10^9$
$10^8$
$10^7$
$10^6$
$10^5$
$10^4$
$10^3$
Buffer
serum
Δlyso serum
serum + OmCI

#### SFig 2

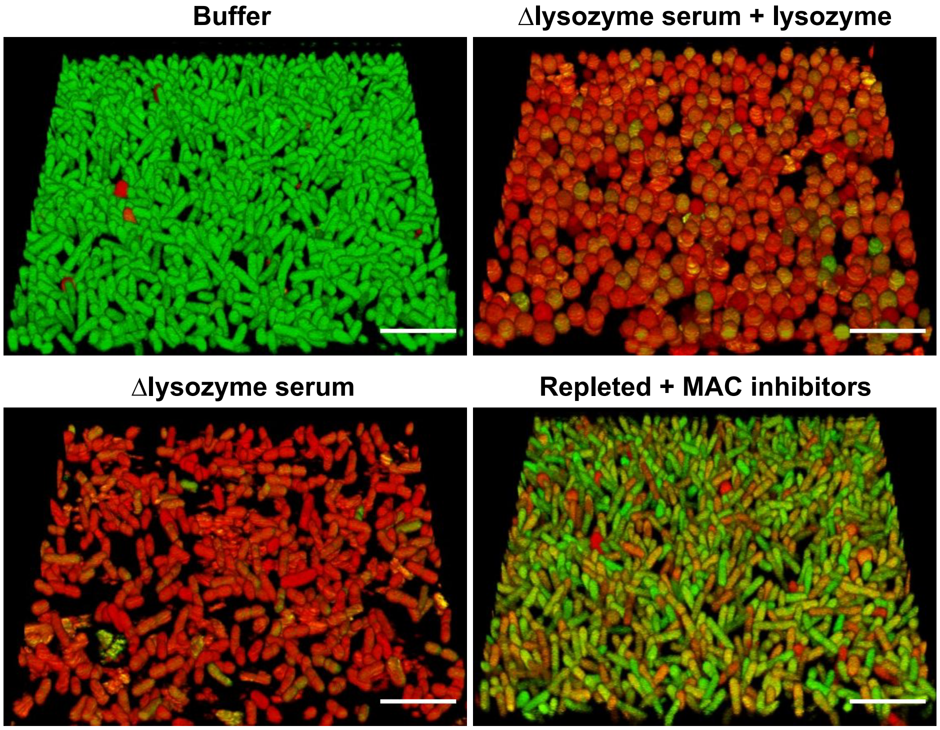
Buffer
Δlysozyme serum + lysozyme
Δlysozyme serum
Repleted + MAC inhibitors

**SFig 3**

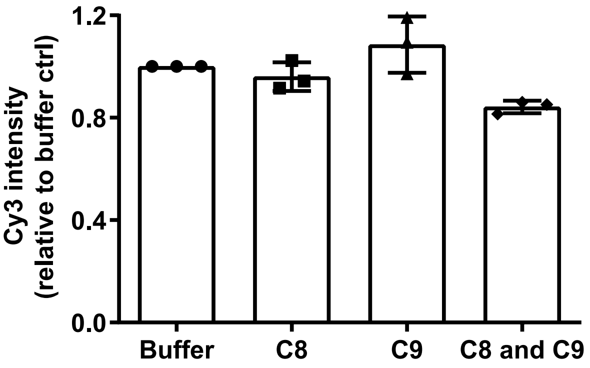

Cy3 intensity
(relative to buffer ctrl)
1.2
0.8
0.4
0.0
Buffer
C8
C9
C8 and C9
